# Physical-aware model accuracy estimation for protein complex using deep learning method

**DOI:** 10.1101/2024.10.31.621211

**Authors:** Haodong Wang, Meng Sun, Lei Xie, Dong Liu, Guijun Zhang

## Abstract

With the breakthrough of AlphaFold2 on monomers, the research focus of structure prediction has shifted to protein complexes, driving the continued development of new methods for multimer structure prediction. Therefore, it is crucial to accurately estimate quality scores for the multimer model independent of the used prediction methods. In this work, we propose a physical-aware deep learning method, DeepUMQA-PA, to evaluate the residue-wise quality of protein complex models. For the input complex model, the residue-based contact area and orientation features were first constructed using Voronoi tessellation, representing the potential physical interactions and hydrophobic properties. Then, the relationship between local residues and the overall complex topology as well as the inter-residue evolutionary information are characterized by geometry-based features, protein language model embedding representation, and knowledge-based statistical potential features. Finally, these features are fed into a fused network architecture employing equivalent graph neural network and ResNet network to estimate residue-wise model accuracy. Experimental results on the CASP15 test set demonstrate that our method outperforms the state-of-the-art method DeepUMQA3 by 3.69% and 3.49% on Pearson and Spearman, respectively. Notably, our method achieved 16.8% and 15.5% improvement in Pearson and Spearman, respectively, for the evaluation of nanobody-antigens. In addition, DeepUMQA-PA achieved better MAE scores than AlphaFold-Multimer and AlphaFold3 self-assessment methods on 43% and 50% of the targets, respectively. All these results suggest that physical-aware information based on the area and orientation of atom-atom and atom-solvent contacts has the potential to capture sequence-structure-quality relationships of proteins, especially in the case of flexible proteins. The DeepUMQA-PA server is freely available at http://zhanglab-bioinf.com/DeepUMQA-PA/.

## 1. Introduction

Protein-protein complexes are central in many crucial biological and cellular processes, which makes their structural elucidation important. With the significant progress made by AlphaFold2 [1] in single-chain structure prediction, the prediction of structures for protein multimers has become the focus of research in the field. Since the structure of protein complexes is the key to understanding its function, methods such as AlphaFold-Multimer [2], DMFold-Multimer [3], AFsample [4], trRosettaX2 [5], and recently released AlphaFold3 [6] have been actively developed to predict the structure of multimers. Nonetheless, challenges remain in predicting structures with weak evolutionary signals, such as nanobody-antigen and antibody-antigen complexes [7]. The CASP15 results show that most successful prediction methods for protein multimers used modifications of the standard AlphaFold, including extensive sampling through variations on MSA construction, the use of multiple seeds, an increased number of recycles and extensive network dropout [7]. It also shows that, at least for now, scoring and ranking the accurate models from many decoys has become a fundamental strategy for improving the accuracy of protein multimers structure prediction. Not surprisingly, estimation of model accuracy (EMA) of multimeric structures has recently received much attention in the field and has been introduced into CASP15 as a new prediction category [8].

Generally, the EMA methods of complex are divided into two categories: multi-model methods and single-model methods. Multi-model methods require multiple models as input and then evaluate the quality of the models using structural alignments and strategies, such as MULTICOM_qa [9], ModFOLDdock [10] and VoroIF-jury [11]. Single-model methods require only a single protein complex structure as input, and do not require additional information to predict model quality, such as VoroIF-GNN [12], DeepUMQA3 [13], et.al. Multi-model methods have significant advantages in specific scenarios such as Critical Assessment of Techniques for Protein Structure Prediction (CASP). However, they rely heavily on the quality of the model pool, which is closely related to the accuracy of structure prediction. By contrast, single-model EMA methods are not restricted by the model pool and can score as well as select models lightly and efficiently. We have observed that the single-model EMA methods outperform the multi-model method, and even the CASP-specific consensus method, on the accuracy estimation track of complex interface contact residues of CASP15 in 2022. In addition, the single-model EMA method can not only evaluate the accuracy independently of structure prediction methods, but also can be a key component of multi-model methods. Naturally, it has become a frontier and research hotspot in the field of protein structure model quality assessment.

Most single-model EMA methods mainly consider the geometric and evolutionary factors of protein structure models and utilize the deep learning network to reveal the relationship between structure features and model quality. In recent two years, our in-house developed DeepUMQA3 [13] extracts feature from three levels of overall geometric topology, intra-chain and inter-chain, and use the improved deep residual neural network to predict the accuracy of interface residues. AlphaFold-Multimer uses the Evoformer module to encode multiple sequence alignment and template information to reflect evolutionary information and decodes the structural coordinates and quality scores in the Structure module [2]. However, it is challenging for the above methods to characterize the solvent effects of the surrounding environment of the protein surface, which is a crucial driver for the protein folding problem and protein-protein interactions [14]. It is worth noting that VoroIF-GNN predicts the contact area accuracy for the complex interface by building a graph to represent the local contact and solvent surface based on Voronoi tessellation [12, 15, 16]. Inspired by concepts of atom-atom and atom-solvent contact areas, given the strong correlation between surface areas and physical interactions [17], it is reasonable to assume that the orientation characteristics of the contact surface may contain the crucial information of the native structure interface. This hypothesis suggests that, based on geometric and evolutionary features, further considering physical-aware information (e.g., the solvent energy characterized by contact surface area and orientation) may reveal more intrinsic relationships between protein structure and model accuracy.

Based on our previously developed DeepUMQA3 protocol, this work proposes a single-model method, DeepUMQA-PA, which is used for model scoring and ranking of multimeric protein structure. We design physical-aware features based on residue contacts to capture the relationship of hydrophobicity and orientation of the interface, while combining topological and protein sequence embedding to describe geometric and evolutionary features. These representations are fed into an equivalent graph neural network (EGNN) [18] coupled with the invariant point attention mechanism (IPA) [1] and a ResNet network to predict the per-residue accuracy estimation for protein multimers. The test results show that the physical-aware, geometric topological, and evolutionary features are complementary, and the use of these features can significantly improve the performance of accuracy estimation for protein complexes.

## 2. Methods

DeepUMQA-PA is a single-model protein complex EMA method, which includes four main parts: data preparation, feature extraction, network architecture, and residue-wise plDDT scores [19]. The schematic diagram of the designed pipeline is illustrated in Fig. 1, and each part of the pipeline will be described in detail below.

**Fig 1.**
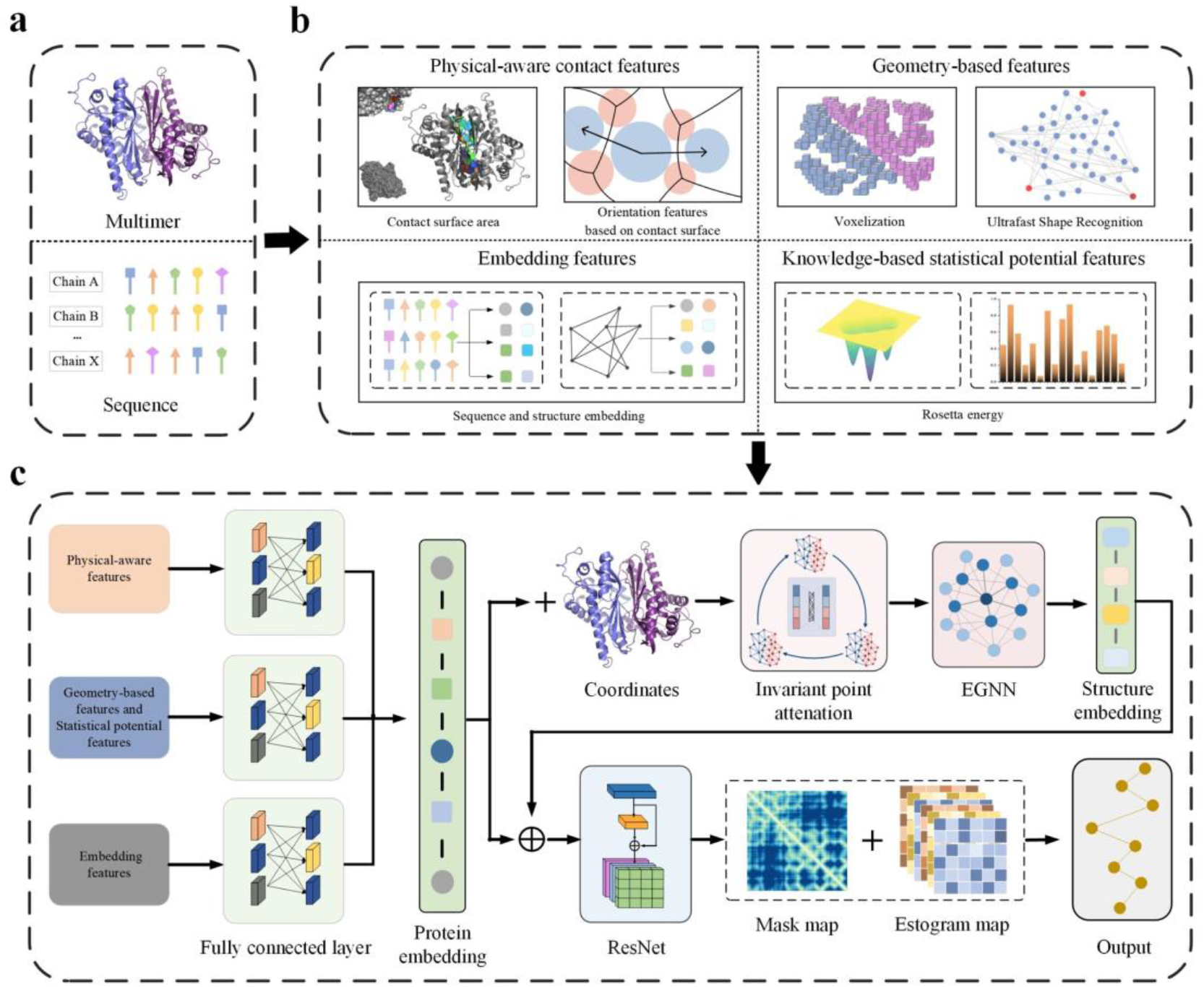
The pipeline of DeepUMQA-PA. **a** Data preparation. Protein complex structure and sequences were taken as input. **b** Features extraction. Physical-aware contact features, geometry-based features, embedding features, and knowledge-based statistical energy features of given protein complex structure are extracted from the input structural and sequence information. **c** Network architecture. The graph neural network was fused with ResNet network with attention mechanisms to estimate residue-wise prediction accuracy.

### 2.1. Feature Extraction

We extract four classes of features from an input protein complex structure: physical-aware contact features (i.e., contact surface area and contact surface-based orientation features), geometry-based features (i.e., ultrafast shape recognition and voxelization features), embedding features (i.e., sequence and structure embedding features), and knowledge-based statistical potential energy features (i.e., Rosetta energy), as shown in Fig. 1B. Details of all these features are available in Table S1 in the supplementary material.

#### 2.1.1 Physical-aware Contact Area feature

In existing literature, most protein complex EMA methods use a distance threshold, such as 5Å or 8Å, between specific atoms (e.g., Cβ and Cα) to define the concept of contact [20-22]. Although this way can effectively characterize the spatial relationships between atoms, it cannot fully reflect the solvent effects of the environment around the protein surface, which are closely related to the atom-atom interactions. Inspired by the concepts of atom-atom and atom-solvent contact area [12, 14-16], we use residue-level contact areas and contact surface-based orientation features to represent the strength of physical interactions between protein surface regions and their surrounding solvents, which maybe provide a new perspective and effective way to evaluate the accuracy of complex structure models.

Given an input complex structure, we first calculate the interatomic contact surface based on the Voronoi tessellation algorithm [15, 16]. For any contact pair formed by atoms *ai* and *aj*, the contact area can be calculated by using the triangulation algorithm [23]. Furthermore, we computed residue-level contact area (RCA) by adding the relevant atom-level contact areas (ACA). Formally, the formula is defined as follows:

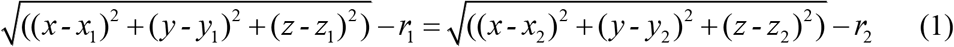

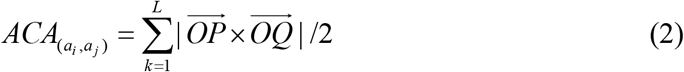

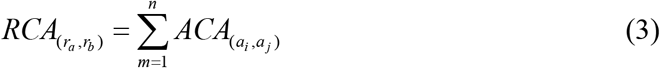

where *r*1 and *r*2 represent the van der Waals radius of atom *ai* and atom *aj* respectively, and the point (*x, y, z*) lies on the contact surface equidistant from the van der Waals spheres of the two atoms (Fig. 2a). *ACA*_(*a*_*i, a j*) represents the contact area between atom *ai* and atom *aj. O, P, Q* are the triangle vertices of the contact surface by triangulation, and *L* represents the total number of triangles. *RCA*_(*ra*_, *r*_*b*_) represents the contact area between residue *ra* and residue *rb. n* represents the total number of atom-level contact surface between residue *ra* and residue *rb*. The schematic diagrams of the residue contact surface and residue-solvent contact surface are shown in Fig. 3.

**Fig 2.**
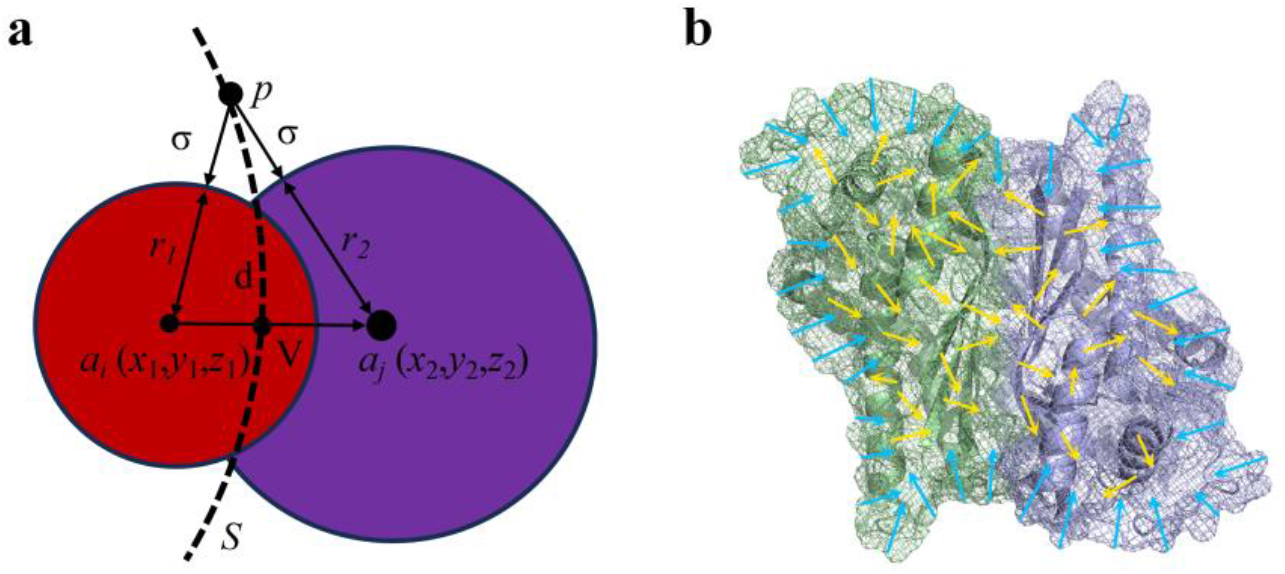
**a** Schematic diagram of atomic contact between atom *ai* (red) and atom *aj* (purple). *r*1 and *r*2 are the van der Waals radius of atom *ai* and atom *aj, d* is the distance between atoms, and *p* is a point on the Voronoi surface, which the surface distances from *p* to atom a and atom b are both *σ*. **b** Schematic diagram of the residue-level contact surface orientation for homodimer 1B5D. The blue arrow represents the contact surface orientation of surface residues, and the yellow arrow represents the contact surface orientation of internal residues.

**Fig 3.**
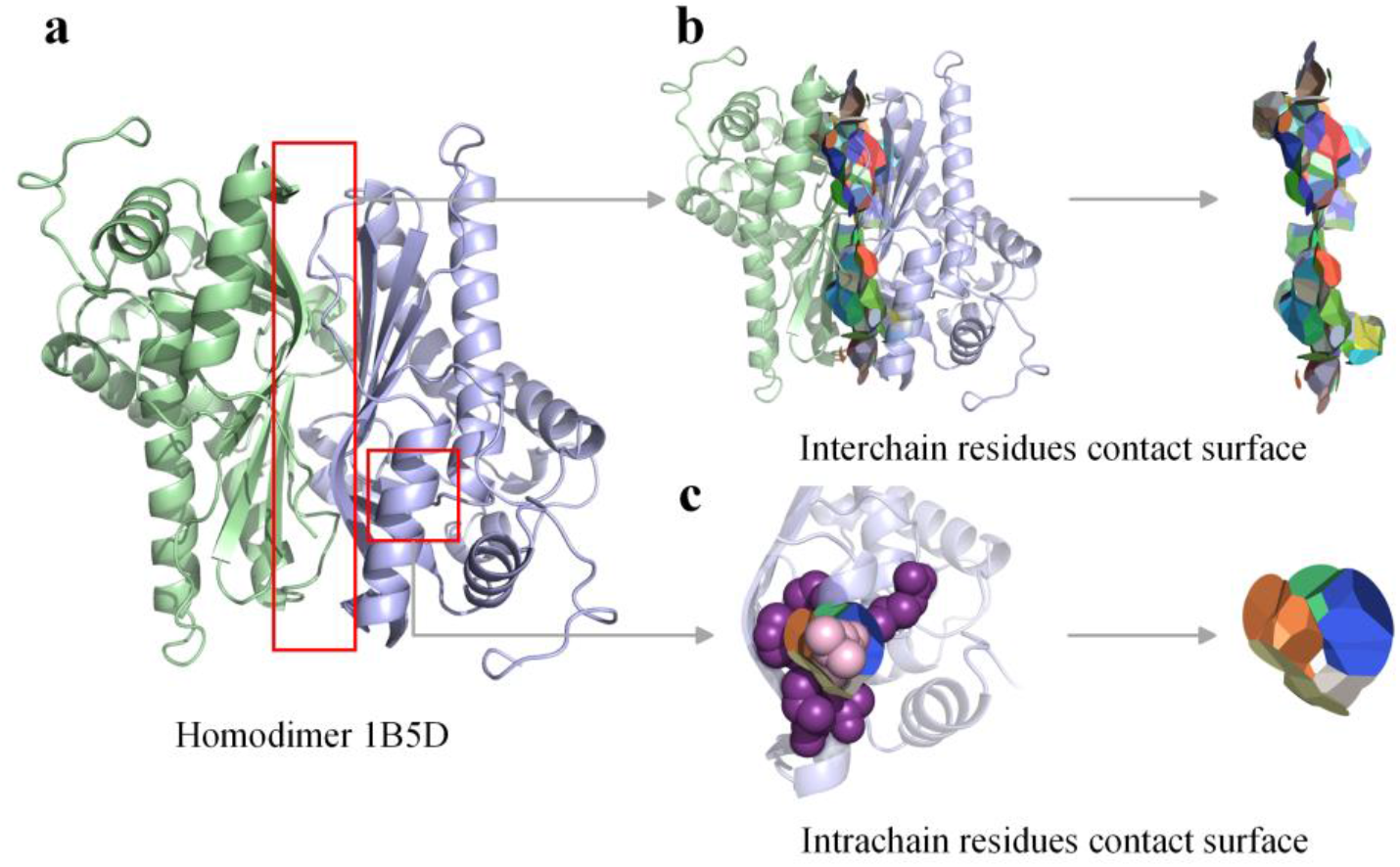
**a** Homodimer 1B5D. **b** Interchain residues contact surface. Random color patches represent the contact surface of different contact residues from interchain. **c** Intrachain residues contact surface. Random color patches represent the contact surface between the pink residue and the other contact residues (purple).

#### 2.1.2 Physical-aware contact orientation feature

To characterize the relationship between protein surface residues and internal residues, we designed residue-level contact surface orientation features, which is the sum of vectors of the contact orientation between a residue and all surrounding contact residues. First, for any two contact atoms *ai* (*x*1, *y*1, *z*1) and *aj* (*x*2, *y*2, *z*2) in a protein, we obtain the contact surface between atoms according to Voronoi tessellation [15]. The atomic contact surface formula in 2.2.1 can be simplified as follows:

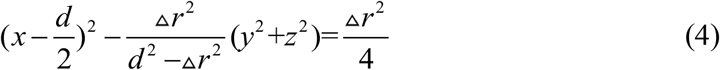

where *r* is the difference in van der Waals radius between atom *ai* and atom *aj, d* is the distance between atoms.

Next, the contact surface *S* between atom *ai* and atom *aj* intersects the connecting line of the two atoms to obtain the reference point *V* (Fig. 2a). We use the support vector 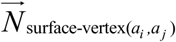 at point *V* as the orientation of the interatomic contact surface. For two residues *ra* and *rb*, the residue-level contact surface orientation 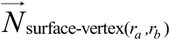 is obtained by summing the support vectors of the relevant atom-level contact surface (Fig. 2b). Formally, the calculation formula is as follows:

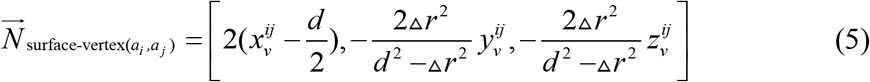

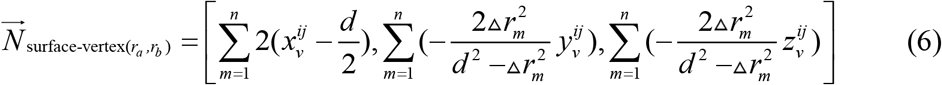

where 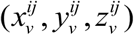 is the coordinate of reference point *V, n* represents the total number of atom-level contact surface between residues *ra* and *r*_*b*_, Δ *rm* is the van der Waals radius difference of the *m*-th atom-level contact surface. More calculation formulas are listed in Table S2 in the supplementary material.

#### 2.1.3 Geometry-based Voxelization and Ultrafast Shape Recognition (USR) features

For protein complex, small conformational changes in local residues could cause a significant impact on inter-chain interactions and the overall structure. The voxelization features project the protein structure into voxelized grids to describe the local structural information of residues [24, 25] while USR [13, 26-29] can quickly capture the topological information of protein structures by using three sets of interatomic distances. Thereby, we use voxelization and overall USR to complementarily characterize the geometric topological relationship between local residues and the overall structure.

#### 2.1.4 Embedding features and statistical potential features

Large Language Models (LLM) capture the evolutionary conservation information of proteins and have been widely used in protein structure prediction, design and function research [30]. The protein language model ESM can quickly and accurately obtain embedding information for structure and sequence [31]. Thereby, we use the high-dimensional embedding of protein structure of backbone atoms in ESM-IF1 [32] (1\**L*\*512) and the high-dimensional embedding of sequence in ESM2 [33] (1\**L*\*1280) to establish the connection between evolutionary conservation information and structural accuracy. In addition, Rosetta energy [34-36] features are also an important part of the input features, using the one-body-terms, the two-body energy terms and the presence of backbone-to-backbone hydrogen bonds features. All these features are normalized and fed to the deep learning neural network.

### 2.2 Network architecture

In this study, we designed a fusion network architecture, where each part is drawn from an equivalent graph neural network (EGNN) [18] coupled with invariant point attention (IPA) [1] and a ResNet network [37] with attention mechanism. The purpose of introducing EGNN network is to process the input structural coordinates, so that we can obtain a structure embedding representation that is closer to the native structure of protein. In this way, by combining feature embeddings from fully connected layers and input into the ResNet network as previously used in DeepUMQA3, we can make more accurate predictions on the distance contact (i.e., Mask) map and the distance error (i.e., Estogram) maps, and finally improve the performance of residue-wise accuracy estimation. The fusion network architecture is shown in Fig. 1C.

For a given protein complex structure model, we first extract four classes of protein features, which were physical-aware contact features, geometry-based features, embedding features, and knowledge-based statistical potential features. These features are firstly input into the fully connected layer [38] to generate a 1\**L*\*128 protein embedding, which is combined with the coordinates of complex model and input into the EGNN network coupled with IPA module. EGNN network is used to iteratively updates of protein atomic coordinates to better approximate the native structure. Then, output of EGNN (i.e., structure embedding) is recombined with the protein embedding and input into the residual network with attention mechanism [39], which consists of a main residual block and two branch residual blocks. Each residual block contains three 2D convolutional layers with different dilation rates, normalization layers and GELU activation function [40]. Finally, we get the mask map thresholded at 15 Å and estogram map with Cβ distance deviations to calculate the residue-wise plDDT score by different branch residual blocks.

### 2.3 Training

To make a fair comparison with DeepUMQA3 and other state-of-the-art methods, the training and validation sets of DeepUMQA3 are used in this work. Meanwhile, we used the CASP15 dataset as test set. Since the training set of DeepUMQA3 was built before CASP15, we can avoid data leakage as much as possible by means of time cutoff. The training and validation datasets contain a total of 7590 targets, each generating approximately 240 models. The ratio of the number of targets in the training and validation datasets is 9:1. Our best network model took 125 hours to train on a single A100. The network Adam optimizer [41] with a learning rate of 0.01 is used, which decays at a rate of 0.05%. During the training process of the network, the performance of the model is optimized by minimizing loss functions. Specifically, the cross-entropy loss function is used to evaluate the estogram map loss; the binary cross entropy loss function is used to evaluate the mask map loss; the root mean square deviation[42] is used to evaluate the coordinate loss and residue-wise plDDT score loss. Loss function is defined as follows:

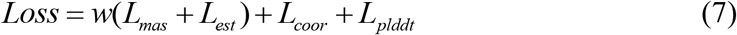

where *w* is the weight that is equal to 0.1, *Lm*as is the mask map loss, *L*_*est*_ is the estogram map loss, *L*_*coor*_ is the coordinate loss, *L*_*plddt*_ is the residue-wise plDDT score loss. lDDT is used to analyze the stability of local regions of proteins. The calculation formula of lDDT is as follows [19]:

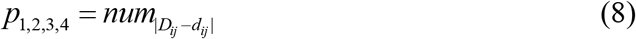

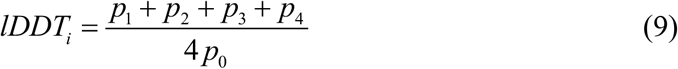

where *p*_0_ represents the number of residues within 15Å of the residue *i* in the reference structure. structure, *Dij* represents the distance between residue *i* and residue *j* in the reference *dij* represents the distance between residue *i* and residue *j* in the prediction model. *p*_1_, *p*_2_, *p*_3_, *p*_4_ respectively represent the number of residues for which the absolute value of | *D*_*ij*_ *-d*_*ij*_ |*<* 0.5 Å, 1 Å, 2 Å, 4 Å. The value range of lDDT is from 0 to 1. The closer the score is to 1, the closer the prediction model is to the reference structure.

## 3. Results and Discussions

In this study, four statistical metrics are used to objectively and fairly analyze the reliability of predictions in complex interface residues. Specifically, we use Pearson [43] and Spearman [44] metrics to measure the correlation between the predicted lDDT score of interface residues and the true lDDT, focusing on evaluating the accuracy of the methods in ranking the models. ROC (AUC) [45] is used to evaluate the ability of the method to distinguish between high-quality and low-quality models. The mean absolute error (MAE) quantifies the deviation between the predicted lDDT scores and the actual values.

### 3.1. Results on the CASP15 test set

We test the performance of DeepUMQA-PA in evaluating the accuracy of interface residues on the CASP15 test set. Due to hardware resource limitations, we compare the performance of the state-of-the-art methods for evaluating local interface accuracy on 7875 models of 30 targets. We find that DeepUMQA-PA has advantages in the reliability of protein complex model scoring and ranking (Fig. 4). The detailed evaluation results are listed in Table S3 of the supplementary material. On average, DeepUMQA-PA outperforms other methods and improves over the top-performing method DeepUMQA3 by 3.69%, 3.49% and 0.48% on Pearson, Spearman and ROC (AUC) of lDDT, respectively (Table 1). In particular, for the five nanobody-antigens targets (H1140-H1144), DeepUMQA-PA significantly improved by 16.8%, 15.5% and 5.1% compared with DeepUMQA3 in the three statistical metrics of Pearson, Spearman and ROC (AUC) based on lDDT, respectively (Table 2). The interaction between antibody and antigen is formed through spatial structural complementarity. The smaller the distance between the two, the greater the interaction force (such as van der Waals force) [46, 47]. This suggests that DeepUMQA-PA may have the potential to accurately assess the local geometry of protein binding sites [48]. This may be attributed to the fact that the introduction of physical-aware features enhances the ability of network to learn specific protein-protein interaction patterns.

**Table 1.** Comparison of the evaluation results of Pearson, Spearman, ROC (AUC) and Z-score on the CASP15 test set with other methods.

| methods | Pearson | Spearman | ROC (AUC) | Z-score |
| --- | --- | --- | --- | --- |
| **DeepUMQA-PA** | **0.608** | **0.574** | **0.765** | **1.356** |
| DeepuMQA3 | 0.587 | 0.554 | 0.761 | 1.332 |
| ModFOLDdockR | 0.476 | 0.440 | 0.686 | 1.144 |
| ModFOLDdockS | 0.425 | 0.393 | 0.663 | 1.072 |
| Venclovas | 0.311 | 0.311 | 0.645 | 0.956 |
| VoroIF | 0.311 | 0.311 | 0.645 | 0.956 |
| FoldEver | 0.294 | 0.288 | 0.631 | 0.922 |
| ModFOLDdock | 0.245 | 0.218 | 0.598 | 0.830 |
| Manifold | 0.130 | 0.115 | 0.546 | 0.669 |

**Table 2.** Comparison of the evaluation results of Pearson, Spearman, ROC (AUC) and Z-score on the nanobody-antigen test set with other methods.

| methods | Pearson | Spearman | ROC (AUC) | Z-score |
| --- | --- | --- | --- | --- |
| **DeepUMQA-PA** | **0.576** | **0.565** | **0.760** | **1.331** |
| DeepuMQA3 | 0.493 | 0.489 | 0.723 | 1.214 |
| ModFOLDdockR | 0.377 | 0.330 | 0.619 | 0.973 |
| ModFOLDdockS | 0.267 | 0.231 | 0.581 | 0.830 |
| FoldEver | 0.206 | 0.210 | 0.595 | 0.803 |
| ModFOLDdock | 0.154 | 0.150 | 0.586 | 0.738 |
| Venclovas | 0.080 | 0.078 | 0.549 | 0.628 |
| VoroIF | 0.080 | 0.078 | 0.549 | 0.628 |
| Manifold | -0.152 | -0.153 | 0.528 | 0.376 |

**Fig 4.**
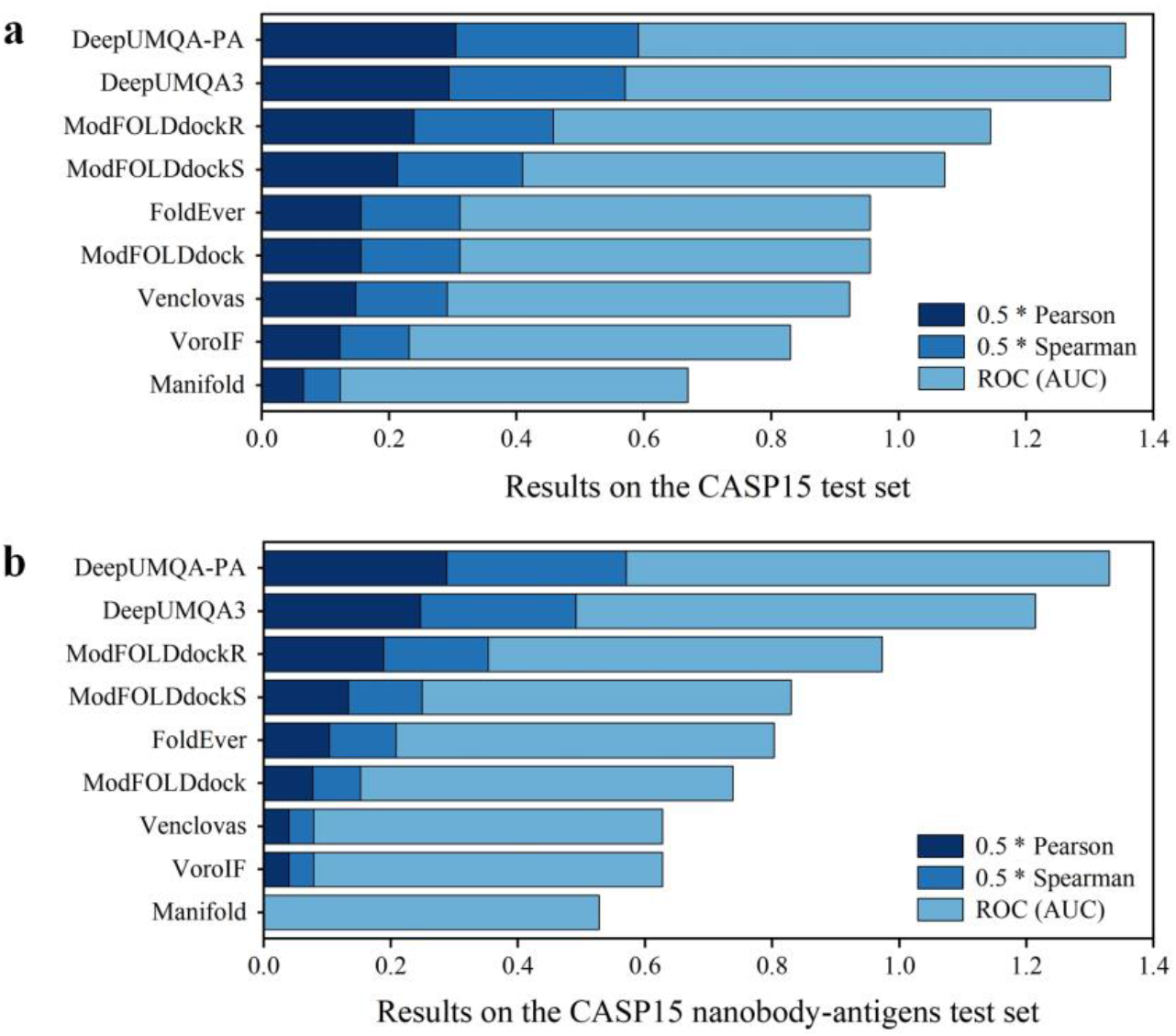
**a** The results of DeepUMQA-PA and other servers on CASP15 test set. **b** The results of DeepUMQA-PA and other servers on CASP15 nanobody-antigens test set.

### 3.2. Ablation studies

In the ablation study, we investigate the impact of physical-aware contact features on the performance of DeepUMQA-PA (Fig. 5). Specifically, we use the same training process to train multiple neural network models with different physical-aware features (i.e., physical-aware orientation, residue-residue contact area and residue-solvent contact area) on the DeepUMQA3 data set, and use 7875 models of 30 targets from CASP15 as the test set. When the physical-aware contact orientation feature is removed from the baseline model DeepUMQA-PA (the full information is used), the Pearson and Spearman of lDDT decrease by 3.49% and 3.22%, respectively. In particular, for the five nanobody-antigen complexes, the Spearman and ROC (AUC) of lDDT decrease by 1.95% and 1.44%, respectively. The difference implies that the introduction of orientation information between residues enables DeepUMQA-PA to consider protein-protein docking and physical interaction mechanisms, which is crucial for identifying binding sites.

**Fig 5.**
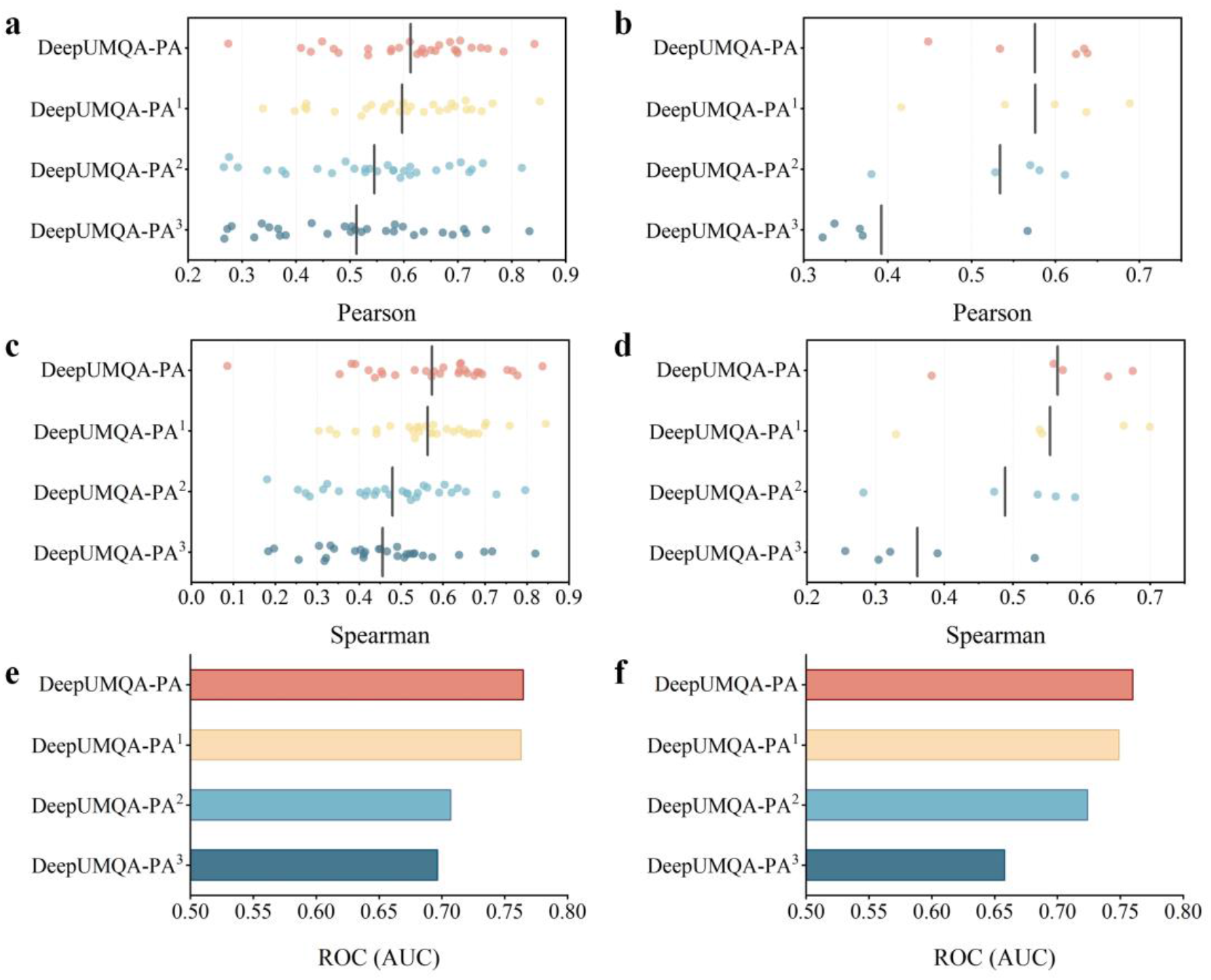
**a c e** Ablation studies of interface residue assessment accuracy on the CASP15 test set. **b d f** Ablation studies of interface residue assessment accuracy on CASP15 nanobody-antigens test set. DeepUMQA-PA is the baseline model that uses all information. DeepUMQA-PA^1^ denotes a version of DeepUMQA-PA without the physical-aware contact orientation feature, and DeepUMQA-PA^2^ additionally removes the physical-aware residue-residue contact area feature. DeepUMQA-PA^3^ represents the model without the physical-aware residue-solvent contact area feature.

We further remove the physical-aware residue-residue contact area feature to investigate the impact of the features on the experimental results. The results shows that the performance of DeepUMQA-PA significantly decrease by 8.29%, 14.48% and 7.32% on the Pearson, Spearman and ROC (AUC) of lDDT respectively. This suggests that the residue-residue contact area feature may play a crucial role in characterizing the physical interactions between interface residues. When the residue-solvent contact area is not used, the performance of the model consistently decreases, with the Pearson correlation decreasing from 0.54 to 0.50 and the Spearman correlation decreasing from 0.47 to 0.45. We also conduct tests on five nanobody-antigen complexes. Notably, the removal of the residue-solvent contact area feature results in a more obvious performance decrease (Pearson: 0.53 to 0.39, Spearman: 0.49 to 0.36, ROC AUC: 0.72 to 0.66). This is mainly because the distribution of solvent molecules near the antibody-antigen binding site affects the energy state of binding, and the residue-solvent contact area can reflect the solvent effects of the environment around the protein surface. In conclusion, the introduction of physical-aware contact features helps DeepUMQA-PA to accurately predict the quality of interface residues.

### 3.3. Comparison with AlphaFold-Multimer and AlphaFold3 self-estimation methods

AlphaFold-Multimer (AFM) and AlphaFold3 (AF3) can not only predict high-precision models but also provide reliable residue-wise confidence estimates. However, for the AF (i.e., AFM and AF3) model, there is still room for improvement in the accuracy of local evaluation. To analyze whether DeepUMQA-PA has the potential to improve evaluation accuracy for AF models with low self-assessment accuracy, we download the AFM prediction models provided by CASP15 and use the AF3 server to generate five models for each target. On average, AFM achieves essentially the same prediction as DeepUMQA-PA on MAE of lDDT for 30 targets (Table S4a). Figure 6a shows the evaluation error comparison between DeepUMQA-PA and AFM on each target. DeepUMQA-PA achieves lower MAE scores than AFM on 43% targets. We further find that DeepUMQA-PA improves more obviously on targets with a higher average MAE predicted by AFM. Especially for the target with MAE > 0.1 between AFM plDDT scores and the true values, the average MAE of DeepUMQA-PA is significantly lower than that of AFM, with the average decrease from 0.169 to 0.140. Similarly, DeepUMQA-PA outperforms AF3 on 50% targets, and improves by 15.17% on these targets (Fig. 6b). These results highlight the synergic intersection between DeepUMQA-PA and AF in assessing local structural accuracy. DeepUMQA-PA provides complementary assessments in region with high uncertainty in AF predictions, which helps to accurately identify low-confidence regions of the prediction model to guide model refinement.

**Fig 6.**
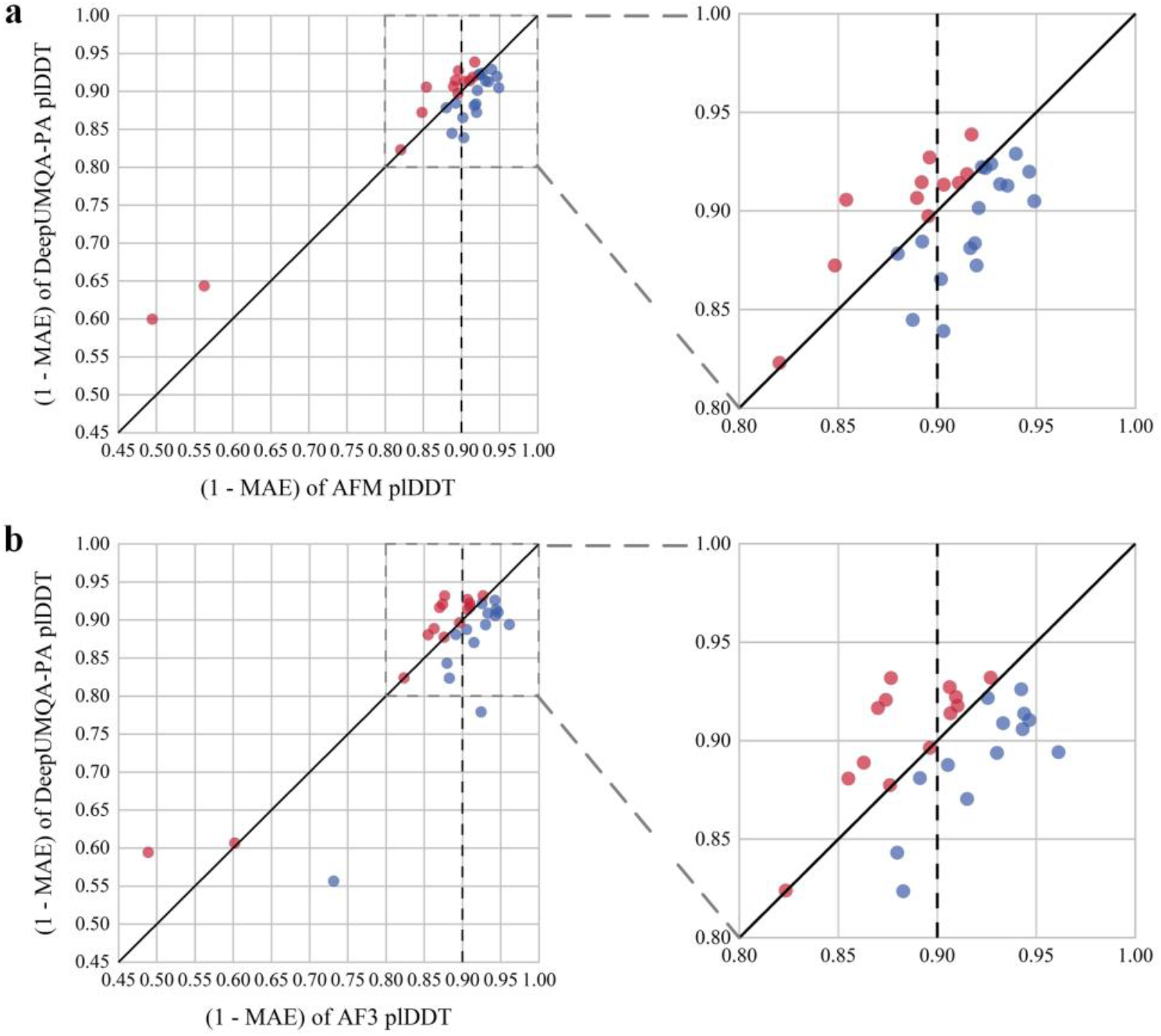
**a** Comparison of DeepUMQA-PA and AlphaFold - Multimer self-assessment accuracy in the mean absolute error (MAE). The models of 30 targets are generated by AFM. **b** Comparison of DeepUMQA-PA and AlphaFold3 self-assessment accuracy in the mean absolute error (MAE). The models of 30 targets are generated by AF3 webserver.

### 3.4. Differences in quality estimation accuracy according to residue location

We divide the protein complex model into core, interface, and surface residues, and observe differences in quality estimation accuracy according to residue location. The classification criteria for residue positions in the complex are as follows: (1) interface residues: Cβ distance between chains≤8 Å (Cα for glycine), (2)core residues: relative solvent-accessible surface area (SASA)≤0.25, (3)surface residues: relative SASA>0.25.

We present the accuracy errors of core, surface, and interface residues predicted by DeepUMQA-PA in Fig. 7. On average, the evaluation accuracy of core residues is 24.79%, 29.33% higher than that of surface and interface residues, respectively (Table S5). Especially for the nanobody antigens (H1140-H1144), the gap in the MAE of lDDT between interface and core residues is even more obvious (core:0.074, surface:0.108, interface:0.174). This may be attributed to the fact that core residues are mainly used to maintain the overall stability of the protein, and their structural patterns are relatively simple and easy to evaluate. In contrast, the structures corresponding to surface and interface residues may undergo conformational changes due to the influence of the surrounding environment or ligands, making the evaluation of these residues more challenging.

**Fig 7.**
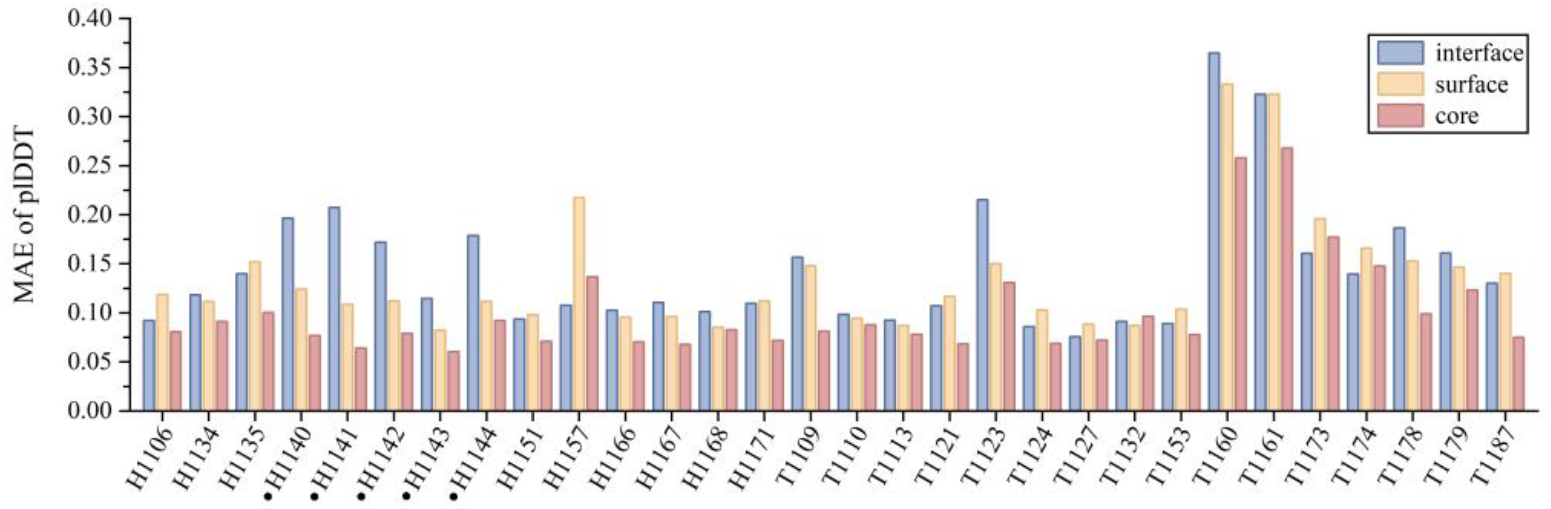
Differences in quality estimation accuracy according to residue location in CASP15 targets. Blue represents interface residues, yellow represents surface residues, and red represents core residues.

## 4. Conclusion

We developed a single-model EMA method for protein complex called DeepUMQA-PA. Based on DeepUMQA3, we further used physical-aware contact surface features (i.e. contact surface area and contact surface-based orientation features) and a fusion network architecture to evaluate the residue-wise model quality. Experimental results demonstrate that our method outperforms state-of-the-art EMA methods, including DeepUMQA3, ModFOLDdockR, ModFOLDdockS, VoroIF, Venclovas, FoldEver, ModFOLDdock and Manifold on 30 protein complex targets in CASP15. Ablation results demonstrate that physical-aware contact surface features can improve the performance of model quality assessment methods. In addition, for the MAE metric, our method is complementary to AlphaFold-Multimer and AlphaFold 3 in terms of local assessment accuracy and has an advantage over it in evaluating low-accuracy models. We further find that it is more challenging for DeepUMQA-PA to evaluate interface residues than core residues and surface residues. With the rapid development of complex structure prediction, model evaluation of protein binding to DNA, RNA and small molecules may be a research hotspot in the future.

## CRediT authorship contribution statement

**Haodong Wang:** Writing - original draft, Writing - review & editing, Conceptualization. **Meng Sun:** Writing - original draft, Writing - review & editing, Conceptualization. **Lei Xie:** Writing - review & editing, Visualization. **Dong Liu:** Writing - review & editing. **Guijun Zhang:** Conceptualization, Supervision, Project administration, Writing - original draft, Writing - review & editing.

## Declaration of Competing Interest

The authors declare that they have no known competing financial interests or personal relationships that could have appeared to influence the work reported in this paper.

## Acknowledgement

This study is supported by the National Key R & D Program of China (2022ZD0115103), the National Nature Science Foundation of China (62173304).

## Supplementary material

**Table S1.** Extracted four classes of features from an input protein complex structure

|  |  |
| --- | --- |
| physical-aware contact features | residue-residue contact surface area [S1]<br>residue-solvent contact surface area [S1]<br>surface-based orientation |
| geometry-based features | ultrafast shape recognition [S2]<br>voxelization feature [S3] |
| embedding features | structure embedding from ESM-IF1 [S4]<br>sequence embedding from ESM2 [S5] |
| knowledge-based statistical potential energy features | one-body-terms (omega, p_aa_pp, fa_dun and rama_prepro)<br>two-body-terms (fa_atr, fa_rep, fa_sol, lk_ball_wtd, fa_elec, hbond_bb_sc and hbond_sc)<br>the presence of backbone-to-backbone hydrogen bonds (hbond_sr_bb and hbond_lr_bb) [S6] |

**Table S2.**
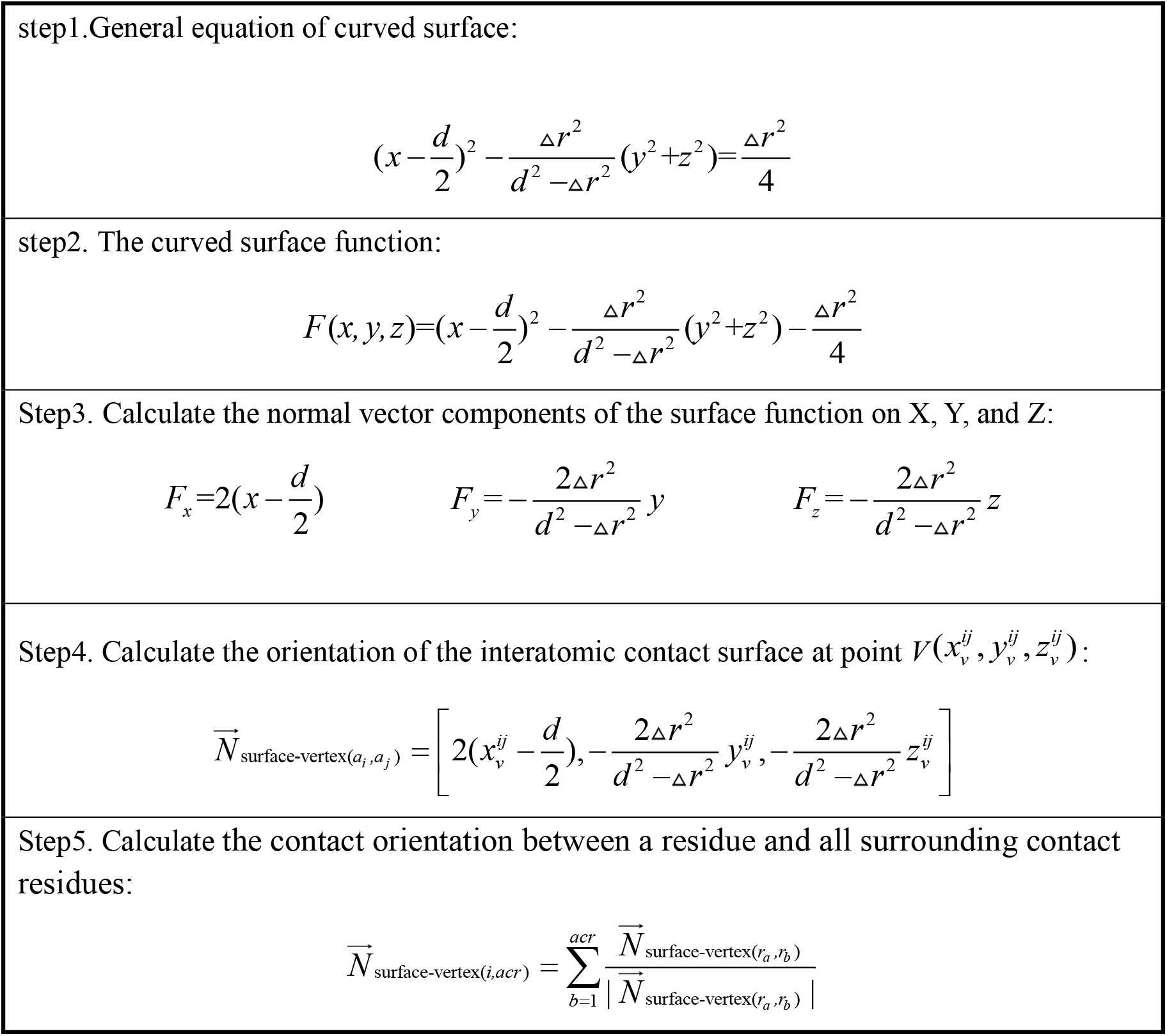
Calculation process of the residue-level contact surface orientation feature

**Table S3.** The performance of DeepUMQA-PA in evaluating the accuracy of interface residues on the CASP15 test set

| **target** | **Pearson** | **Spearman** | **AUC** |
| --- | --- | --- | --- |
| H1106 | 0.665 | 0.600 | 0.791 |
| H1134 | 0.725 | 0.753 | 0.858 |
| H1135 | 0.658 | 0.637 | 0.774 |
| H1140 | 0.534 | 0.559 | 0.785 |
| H1141 | 0.448 | 0.382 | 0.667 |
| H1142 | 0.638 | 0.674 | 0.843 |
| H1143 | 0.634 | 0.639 | 0.806 |
| H1144 | 0.624 | 0.572 | 0.699 |
| H1151 | 0.743 | 0.652 | 0.767 |
| H1157 | 0.591 | 0.532 | 0.717 |
| H1166 | 0.699 | 0.777 | 0.878 |
| H1167 | 0.470 | 0.578 | 0.772 |
| H1168 | 0.576 | 0.593 | 0.766 |
| T1109 | 0.409 | 0.422 | 0.728 |
| T1110 | 0.785 | 0.456 | 0.645 |
| T1113 | 0.698 | 0.658 | 0.766 |
| T1121 | 0.631 | 0.693 | 0.819 |
| T1123 | 0.755 | 0.766 | 0.914 |
| T1124 | 0.577 | 0.452 | 0.695 |
| T1127 | 0.611 | 0.390 | 0.678 |
| T1132 | 0.534 | 0.437 | 0.662 |
| T1153 | 0.650 | 0.686 | 0.863 |
| T1160 | 0.274 | 0.086 | 0.505 |
| T1161 | 0.427 | 0.353 | 0.580 |
| H1171 | 0.490 | 0.573 | 0.808 |
| T1173 | 0.842 | 0.837 | 0.887 |
| T1174 | 0.704 | 0.642 | 0.769 |
| T1178 | 0.479 | 0.486 | 0.776 |
| T1179 | 0.685 | 0.640 | 0.921 |
| T1187 | 0.694 | 0.681 | 0.804 |
| average | 0.608 | 0.574 | 0.765 |

**Table S4a.**
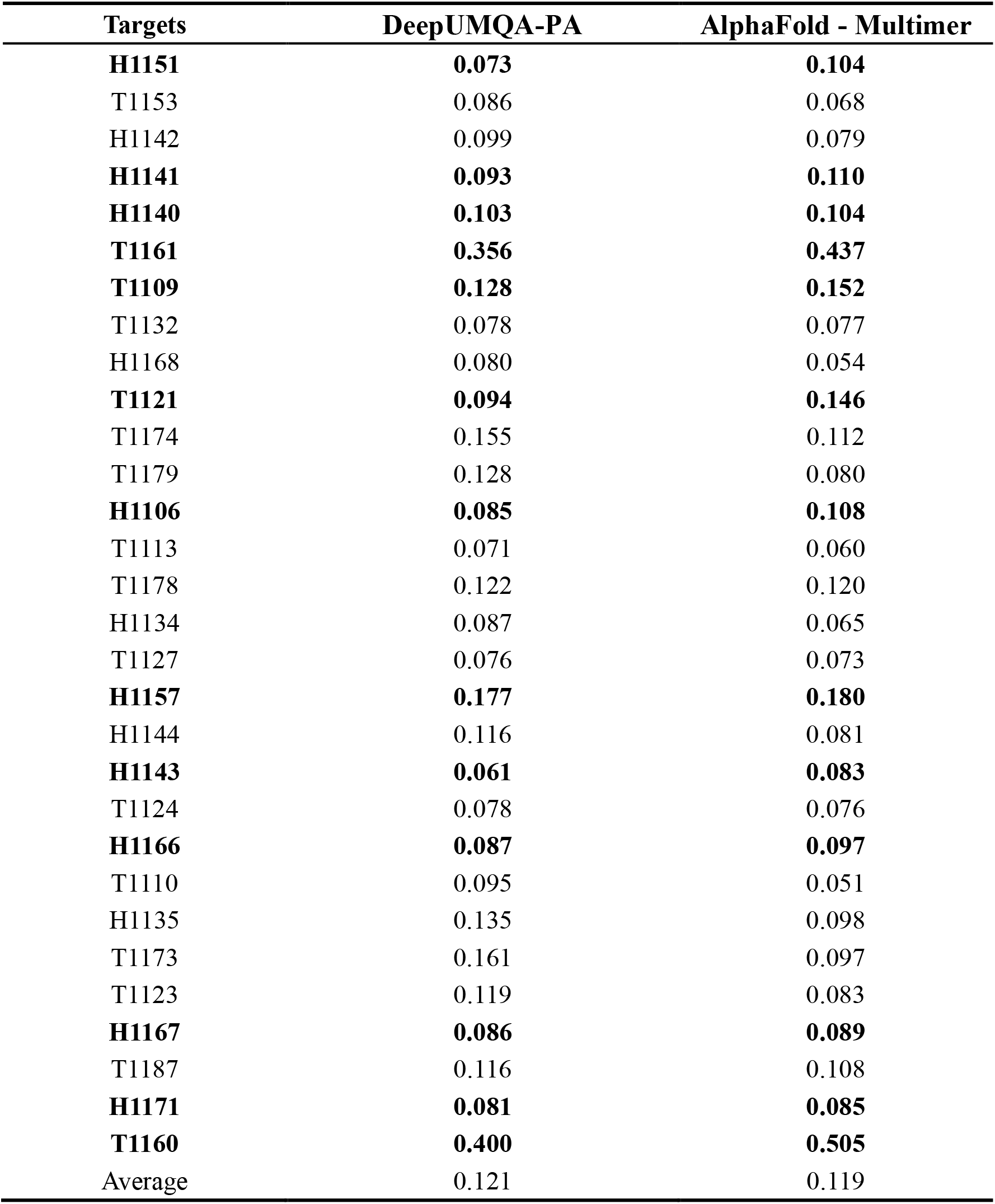
Comparison of DeepUMQA-PA and AlphaFold - Multimer self-assessment accuracy in the mean absolute error (MAE).

| **Targets** | **DeepUMQA-PA** | **AlphaFold - Multimer** |
| --- | --- | --- |
| **H1151** | **0.073** | **0.104** |
| T1153 | 0.086 | 0.068 |
| H1142 | 0.099 | 0.079 |
| **H1141** | **0.093** | **0.110** |
| **H1140** | **0.103** | **0.104** |
| **T1161** | **0.356** | **0.437** |
| **T1109** | **0.128** | **0.152** |
| T1132 | 0.078 | 0.077 |
| H1168 | 0.080 | 0.054 |
| **T1121** | **0.094** | **0.146** |
| T1174 | 0.155 | 0.112 |
| T1179 | 0.128 | 0.080 |
| **H1106** | **0.085** | **0.108** |
| T1113 | 0.071 | 0.060 |
| T1178 | 0.122 | 0.120 |
| H1134 | 0.087 | 0.065 |
| T1127 | 0.076 | 0.073 |
| **H1157** | **0.177** | **0.180** |
| H1144 | 0.116 | 0.081 |
| **H1143** | **0.061** | **0.083** |
| T1124 | 0.078 | 0.076 |
| **H1166** | **0.087** | **0.097** |
| T1110 | 0.095 | 0.051 |
| H1135 | 0.135 | 0.098 |
| T1173 | 0.161 | 0.097 |
| T1123 | 0.119 | 0.083 |
| **H1167** | **0.086** | **0.089** |
| T1187 | 0.116 | 0.108 |
| **H1171** | **0.081** | **0.085** |
| **T1160** | **0.400** | **0.505** |
| Average | 0.121 | 0.119 |

**Table S4b.** Comparison of DeepUMQA-PA and AlphaFold3 self-assessment accuracy in the mean absolute error (MAE).

| **Targets** | **DeepUMQA-PA** | **AlphaFold3** |
| --- | --- | --- |
| **H1151** | **0.073** | **0.094** |
| T1153 | 0.091 | 0.067 |
| H1142 | 0.112 | 0.095 |
| **H1141** | **0.104** | **0.104** |
| **H1140** | **0.068** | **0.123** |
| **T1161** | **0.393** | **0.398** |
| **T1109** | **0.123** | **0.124** |
| T1132 | 0.106 | 0.039 |
| H1168 | 0.086 | 0.056 |
| **T1121** | **0.079** | **0.126** |
| T1174 | 0.157 | 0.120 |
| T1179 | 0.221 | 0.076 |
| **H1106** | **0.083** | **0.130** |
| T1113 | 0.074 | 0.058 |
| **T1178** | **0.119** | **0.145** |
| H1134 | 0.089 | 0.053 |
| **T1127** | **0.078** | **0.091** |
| **H1157** | **0.176** | **0.177** |
| H1144 | 0.106 | 0.070 |
| **H1143** | **0.111** | **0.137** |
| **T1124** | **0.086** | **0.093** |
| **H1166** | **0.082** | **0.090** |
| T1110 | 0.094 | 0.057 |
| H1135 | 0.130 | 0.085 |
| T1173 | 0.443 | 0.268 |
| T1123 | 0.176 | 0.117 |
| H1167 | 0.078 | 0.074 |
| T1187 | 0.119 | 0.109 |
| **H1171** | **0.068** | **0.073** |
| **T1160** | **0.405** | **0.511** |
| Average | 0.138 | 0.125 |

**Table S5.** Differences in quality estimation accuracy according to residue location in CASP15 targets. (DeepUMQA-PA)

| **Targets** | **Interface** | **Surface** | **Core** |
| --- | --- | --- | --- |
| H1144 | 0.179 | 0.112 | 0.092 |
| T1161 | 0.323 | 0.323 | 0.268 |
| T1113 | 0.092 | 0.087 | 0.078 |
| H1167 | 0.111 | 0.096 | 0.068 |
| H1140 | 0.197 | 0.124 | 0.077 |
| H1151 | 0.094 | 0.098 | 0.071 |
| H1166 | 0.103 | 0.095 | 0.070 |
| T1127 | 0.076 | 0.088 | 0.072 |
| H1157 | 0.108 | 0.218 | 0.137 |
| H1142 | 0.172 | 0.112 | 0.079 |
| T1132 | 0.091 | 0.087 | 0.097 |
| T1178 | 0.186 | 0.153 | 0.099 |
| T1160 | 0.365 | 0.333 | 0.258 |
| H1168 | 0.101 | 0.085 | 0.083 |
| T1153 | 0.089 | 0.104 | 0.078 |
| H1106 | 0.092 | 0.119 | 0.081 |
| T1121 | 0.107 | 0.117 | 0.068 |
| T1179 | 0.161 | 0.147 | 0.123 |
| T1187 | 0.130 | 0.140 | 0.075 |
| T1174 | 0.140 | 0.166 | 0.148 |
| H1171 | 0.110 | 0.112 | 0.072 |
| H1143 | 0.114 | 0.083 | 0.060 |
| T1109 | 0.157 | 0.148 | 0.081 |
| H1134 | 0.118 | 0.111 | 0.091 |
| H1141 | 0.207 | 0.109 | 0.064 |
| T1173 | 0.161 | 0.196 | 0.177 |
| T1124 | 0.086 | 0.103 | 0.069 |
| T1110 | 0.099 | 0.094 | 0.088 |
| T1123 | 0.215 | 0.150 | 0.131 |
| H1135 | 0.140 | 0.152 | 0.100 |
| Average | 0.144 | 0.135 | 0.102 |

## Notes

### Competing Interest Statement

The authors have declared no competing interest.

